# Prophage-marine diazotroph interplay shapes both biofilm structure and nitrogen release

**DOI:** 10.1101/2025.07.17.665314

**Authors:** Louise Mahoudeau, Pauline Crétin, Aurélie Joublin-Delavat, Sophie Rodrigues, Clara Guillouche, Isabelle Louvet, Nadège Bienvenu, Claire Geslin, Gabriel Dulaquais, Jean-François Maguer, François Delavat

## Abstract

Marine environments are frequently oligotrophic, characterized by low amount of bioassimilable Nitrogen sources. At the global scale, the microbial fixation of N_₂_, or diazotrophy, represents the primary source of fixed nitrogen in pelagic marine ecosystems, playing a key role in supporting primary production and driving the export of organic matter to the deep ocean. However, given the high energetic cost of N_₂_ fixation, the active release of fixed nitrogen by diazotrophs appears counterintuitive, suggesting the existence of alternative, passive release pathways, that remain understudied to date. Here, we show that the marine Non Cyanobacterial Diazotroph *Vibrio diazotrophicus* is endowed with a prophage belonging to the *Myoviridae* family, whose expression is induced under anoxic and biofilm-forming conditions. We demonstrate that this prophage can spontaneously excise from the genome of its host and that it forms intact and infective phage particles. Moreover, phage-mediated host cell lysis leads to increased biofilm production as compared to a prophage-free derivative mutant, and to increased release of Dissolved Organic Carbon and of ammonium. Altogether, we provide evidences that viruses may play a previously unrecognized role in oceanic ecosystem dynamics by structuring micro-habitats suitable for diazotrophy and by contributing to the recycling of (in-)organic matter.

**Importance:** Diazotrophs are key players in ocean functioning by providing fixed nitrogen to ecosystems and fueling primary production. However, from a physiological point-of-view, the active release of nitrogenous compounds by diazotrophs is paradoxical, since they would invest in an energy-intensive process and supply nutrient to non-sibling cells, with the risk of being outcompeted. Therefore, alternative ways leading to the release of fixed nitrogen must exist. Here, we show that the marine Non Cyanobacterial Diazotroph *Vibrio diazotrophicus* possesses one prophage, whose activation leads to cell death, to increased biofilm production and to the release of Dissolved Organic Compounds and of ammonium. Taken together, our results provide evidences that marine phage-diazotroph interplay leads to the creation of micro-habitats suitable for diazotrophy like biofilm and to nutrient cycling, and contribute to better understanding of the role of viruses in marine ecosystems.

## Introduction

The process of dinitrogen (N_2_) fixation, or diazotrophy, is restricted to a small part of polyphyletic prokaryotes known as diazotrophs. In marine environments, diazotrophy is now recognized as a key process in the biogeochemical cycles of nitrogen (N) but also of carbon (C), since it represents the main external source of nitrogen for the upper ocean that stimulates the biological pump, a mechanism that allows a large part of the carbon dioxide emitted into the atmosphere to be absorbed and then exported to the ocean seafloor. The key enzymatic complex in the process is the nitrogenase, which converts N_2_ into NH_3_. The nitrogenase encoding genes are thought to have appeared early in evolution (1), before the Great Oxidation Event (2). Nowadays, this enzymatic complex remains highly sensitive to dioxygen (O_2_) (3), and this constraint represents a major metabolic conflict, since many diazotrophs are also aerobes, performing aerobic respiration to fuel the high ATP-demanding nitrogenase. On top of this constraint, many cyanobacterial diazotrophs are also photoautotrophs, releasing O_2_ as a by-product of oxygenic photosynthesis. To face this metabolic challenge, known diazotrophs have developed various strategies, including spatial separation of diazotrophy in a subpopulation of differentiated cells (4), temporal separation with diazotrophy occurring during the night, *i.e.* without oxygenic photosynthesis, hyper-respiration (5, 6) or synthesis of hopanoid lipids to limit O_2_ diffusion (7). Alternatively, some diazotrophs seek more appropriate conditions, by living inside plant nodules, which create micro-oxic environments suitable for diazotrophy.

Since decades, free Cyanobacteria, particularly those belonging to the genus *Trichodesmium*, are considered to be responsible for much of N_2_-fixation in the oceans (8). However, more recent findings have greatly expanded our understanding of marine N_2_-fixation globally. Indeed, studies based on the amplification of *nifH–*a recognized marker of N_2_-fixation*–* (9, 10) along with whole genomes assembly from Tara Ocean expeditions (11) have shown the widespread occurrence of Non-Cyanobacterial Diazotrophs (NCDs) in the oceans., Remarkably, NCDs were sometimes found in greater abundance than Cyanobacteria, accounting for up to 92.6% of the diazotrophs in the collected metagenomes from the 0.8 μm-2000 μm size fraction (11). These results suggest that NCDs may play a major role in the marine N_2_-fixation activity. Until now, the strategies deployed by these NCDs during N_2_-fixation remain understudied. We recently demonstrated that the marine diazotroph *Vibrio diazotrophicus* is able to utilize a wide variety of organic and inorganic sources upon shifting to diazotrophic conditions and to hyper-respirate to decrease O_2_ tension. Moreover, we showed that the strain modulates the proportion of nitrogenase-expressing cells during diazotrophic growth in an ammonium-dependent manner, and that this phenotypic heterogeneity might be a conserved trait within marine NCDs (12). In addition, marine NCDs are frequently found in large size fractions of the water column, suggesting cell aggregation and/or colonization of large particles (11). This aggregate/biofilm production may be a strategy to provide low-O_2_ micro-environments required for N_2_-fixation. Supporting this hypothesis, the marine diazotroph *Pseudomonas stutzeri* BAL361 produces cell aggregates under oxic growth with N_2_ as sole nitrogen source (13) and *V. diazotrophicus* NS1 increases biofilm production under Soluble Reactive Nitrogen (SRN) limited conditions (14).

Irrespective of the strategies deployed by diazotrophs, the diazotrophically derived nitrogen is subsequently assimilated by the N_2_-fixing-cell to create new biomass (15). Nevertheless, a fraction of this nitrogen can be released extra-cellularly, potentially fuelling the broader (photoautotrophic) plankton community. This release of organic-and/or inorganic nitrogen-containing molecules might involve dedicated transporters. However, because diazotrophs would invest energy in N_2_-fixation to (in)directly feed non-sibling cells, with a risk of being outcompeted, an alternative way to release the fixed nitrogen may occur, but this has not been investigated. One such alternative might be viruses, whose lytic activity may play fundamental but still understudied role in aquatic food webs (16).

In this study, we demonstrate that biofilm production and the release of carbon and nitrogen to the ecosystem by the marine diazotroph *V. diazotrophicus* at least partly rely on the activation of a prophage. We provide evidences that viruses may play a previously unrecognized role in oceanic ecosystem functioning by structuring micro-habitats suitable for diazotrophs and by the recycling of (in-)organic matter.

## Results

### V. diazotrophicus engages an intensive transcriptomic remodeling to anoxia and in biofilm

*V. diazotrophicus* NS1 was recently shown to produce a thicker biofilm under SRN-limiting conditions (14), while the presence of O_2_ is known to inhibit the nitrogenase. To have a global overview of the adaptive response of this facultative anaerobe when grown under anoxic conditions (MDV anoxia) or within a biofilm (MDV biofilm), we first sought to perform a global transcriptomic approach. Each condition was compared to the reference condition in which cells were grown in liquid culture under oxic conditions (MDV), the MDV medium being a SRN-limited medium. A total of 1540 and 1883 genes were significantly differentially expressed in the comparison of MDV vs MDV anoxia and MDV vs MDV biofilm (**Fig. 1, Fig. S1** and **Table S4**), corresponding to a differential expression of 35-43% of the entire genome (4431 genes). When comparing MDV and MDV in anoxia, 750 genes (49% of the Differentially Expressed Genes DEGs) were significantly upregulated in anoxia while 790 genes (51% of the DEGs) were downregulated (**Table S1** and **S4**). Similar proportions were found when comparing MDV with MDV in biofilm, with 910 genes (48% DEGs) being overexpressed in biofilm and 973 (52% DEGs) being under-expressed in biofilm (**Table S4** and **S1**).

**Fig. 1.**
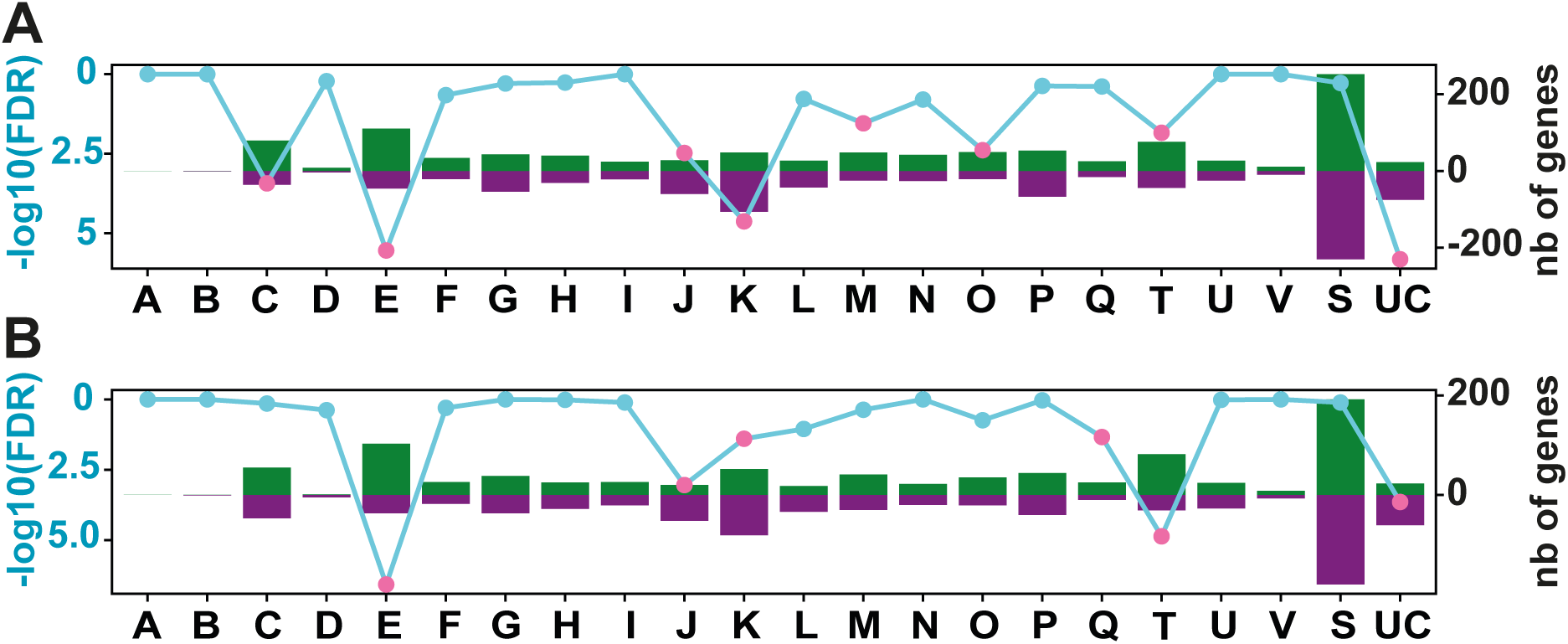
Global transcriptomic response of *V. diazotrophicus* when grown in biofilm (A) or in anoxia (B) as compared to liquid aerobic growth. The vertical bars represent the number of upregulated (green) and downregulated (purple) genes in each COG category. The secondary y-axis (left) shows the statistical significance of the Fisher exact test, represented as - log10(FDR). A threshold of significance (-log10(FDR) ≥ 1.30) is indicated by pink dots, while non-significant values are marked in cyan. COG letters correspond to A: RNA processing and modification. C: Energy production and conversion. D: Cell cycle control, cell division and chromosome distribution. E: Amino acid transport and metabolism. F: Nucleotide transport and metabolism. G: Carbohydrate transport and metabolism. H: Transport and metabolism of coenzymes. I: Lipid transport and metabolism. J: Ribosome translation, structure and biogenesis. K: Transcription. L: Replication, recombination and repair. M: Cell wall, membrane and envelope biogenesis. N: Cell motility. O: Post-translational modification, protein turnover and chaperones. P: Transport and metabolism of inorganic ions. Q: Biosynthesis, transport and catabolism of secondary metabolites. S: Function unknown. T: Signal transduction mechanisms. U: Intracellular traffic, secretion and vesicular transport. UC: Unknown genes. V: Defence mechanisms.

In order to further investigate the metabolic remodeling of this strain to anoxia and biofilm conditions, DEGs were classified by functional classes (COGs, **Fig. 1** and **S1**). In total, between 20.5 and 31,6% of the DEGs were not categorized (either category S “Unknown Function” or UC “Unclassified”, **Fig. S1**).

We subsequently performed a Fisher’s exact overrepresentation test, which determines significantly enriched categories. Interestingly, genes belonging to “Unclassified” category, representing 98 and 94 genes in the two sets of conditions (**Fig. 1**), were significantly enriched, which suggests that many of these unknown genes might be involved in the growth in these particular lifestyles (biofilm and anoxia). The switch from growth under oxic to anoxic conditions (**Fig. 1B**) revealed that *V. diazotrophicus* genes involved in amino acid transport and metabolism (COG E) and signal transduction mechanisms (COG T) are mostly downregulated. In contrast, category J (Translation, ribosomal structure and biogenesis) is significantly up-regulated in anoxia, reflecting intense DNA-to-protein activity. This activity was also observed when comparing MDV with MDV biofilm (**Fig. 1A**), genes from categories J and K being also mostly overexpressed. In contrast, genes from cells grown in MDV biofilm and categorized in COG C (energy production and conversion), E (amino acid transport and metabolism) M (cell wall/membrane/envelope biogenesis), O (posttranslational modification, protein turnover, chaperones) and T (signal transduction) were mostly downregulated, revealing intense cellular reprogramming upon growth in biofilm.

A key parameter that differs between aerobic MDV growth and growth under biofilm or anoxic conditions is the O_2_ tension. Consistent with this parameter change, the *bo* cytochrome (*cyoABCD* genes, BBJY01_570181 to 570184 according to MAGE nomenclature), which operates at high O_2_ tension; was downregulated in both biofilm and anoxic conditions, while the fumarate reductase (*frdABCD* genes, BBJY01_490101 to 490104) were upregulated under anoxia. This latter system allows anaerobic respiration, suggesting that *V. diazotrophicus* is able to use fumarate as alternative terminal electron acceptor.

Strikingly, the entire *nif* cluster, composed of four regions and spanning 100 kb of the *V. diazotrophicus* NS1 genome, was downregulated (from 2- to 30-fold) under anoxia and when grown in biofilm (**Fig. S2**). This includes the *nifHDK* genes encoding the key enzymatic nitrogenase complex, and suggests that diazotrophic activity is repressed under such conditions.

### RNAseq analysis reveals the existence of a prophage region in V. diazotrophicus

In order to have an overview of genomic regions which revealed similarly regulated (indicating operon organization and/or co-regulation of genes), the list of differentially expressed genes was sorted according to their gene order, from gene BBJY01_10001 to BBJY01_570386 according to MAGE nomenclature (https://mage.genoscope.cns.fr/microscope/mage/index.php). This clustering approach led to the discovery of a 40 kb-region, that revealed similarly overexpressed (from 3- to 40-fold) under both anoxic and biofilm conditions (**Table S5** et **Fig. S3**). This region comprised genes BBJY01_510063 to BBJY01_510112, mostly encoding uncharacterized proteins, but with some genes having low-but-significant similarities with phage genes (**Fig. 2**). Interestingly, this 40 kb-region encompassed a potential prophage region identified by the Phigaro tool of the MAGE platform. Phigaro suggested a 20,396 bp long prophage region composed of 24 genes (BBJ001_510081 to BBJ001_510104 using to MAGE annotation). Given the discovery of a potentially co-regulated 40 kb region by transcriptomic analysis and the fact that this region encompassed a potential prophage region, we hypothesized that the actual prophage region was longer than the one initially discovered by Phigaro and was in fact 40 kb-long. This hypothesis was further reinforced by the detection of two tRNA genes flanking this region (**Fig. 3A**), tRNA genes being known hotspots for phages-and other mobile genetic elements-insertion (17, 18). Moreover, a perfect 68 bp duplicated sequence flanked this 40 kb-region and overlapped with the end of the Pro-tRNA gene sequence (found in the left end of the region, **Fig. 3A**). This potential 40 kb-prophage region was considered as the potential prophage region and its corresponding DNA sequence was used for further analyses. We subsequently analyzed the potential prophage genome with PhageScope (19). Analysis revealed that the prophage belongs to the *Myoviridae* family, characterized by their icosahedral head and by their contractile tail sheath (20). This prophage region comprised the genes encoding an integrase, required for phage DNA site-specific integration, an excisionase required for phage DNA excision and a CII protein, responsible for the lysogenic/lytic phase switch (**Fig. 2**). Genes encoding the structure of the viral particle were also present, as well as a gene encoding a potential holin required for cell lysis. Taken together, this region comprised all genes required to produce an active prophage, that we named Vdi_1.

**Fig. 2.**
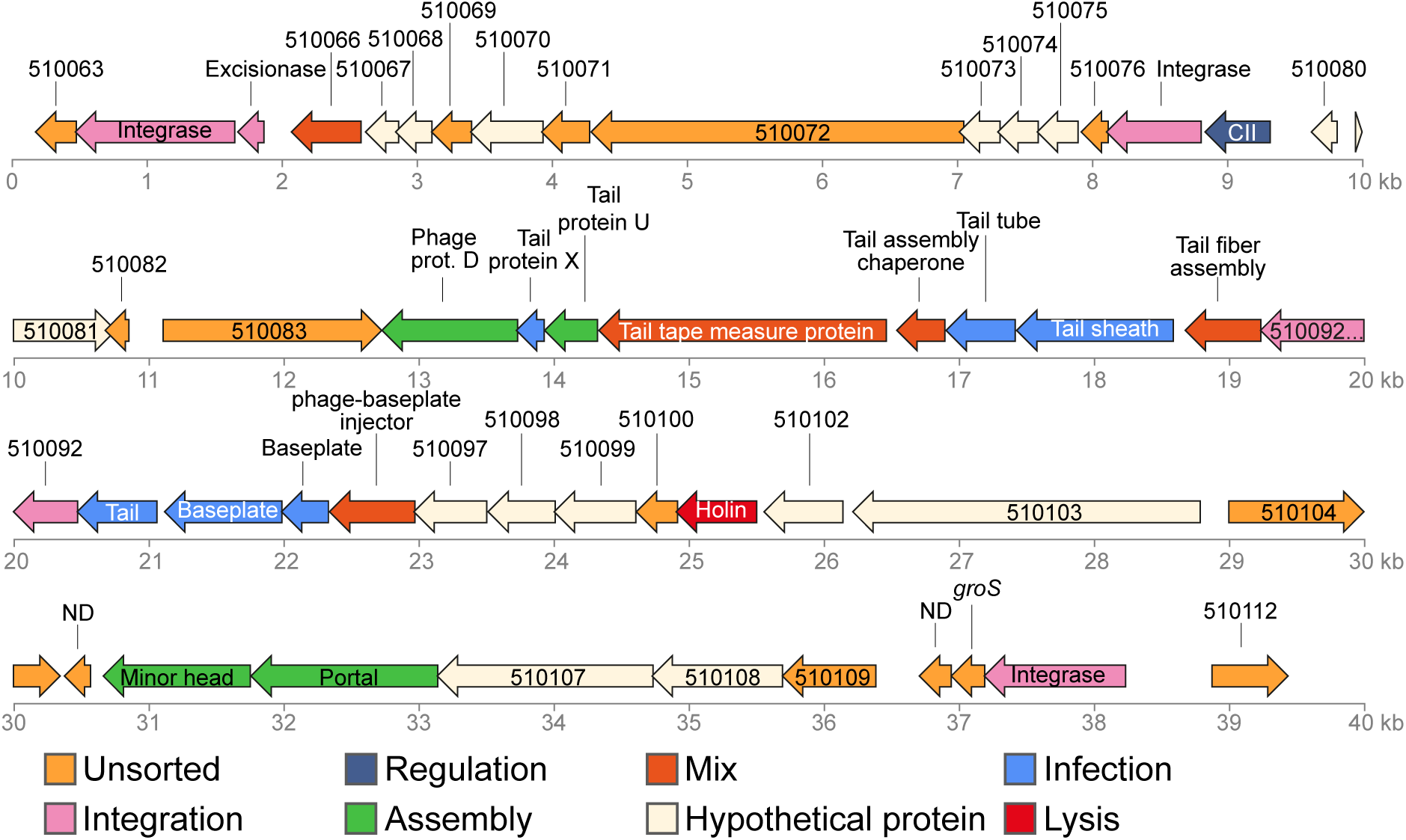
Vdi_1 prophage genes. Every color corresponds to a putative function, according to PhageScope {Wang, 2024 #1478} and Virfam {Lopes, 2014 #1517}.

**Fig. 3.**
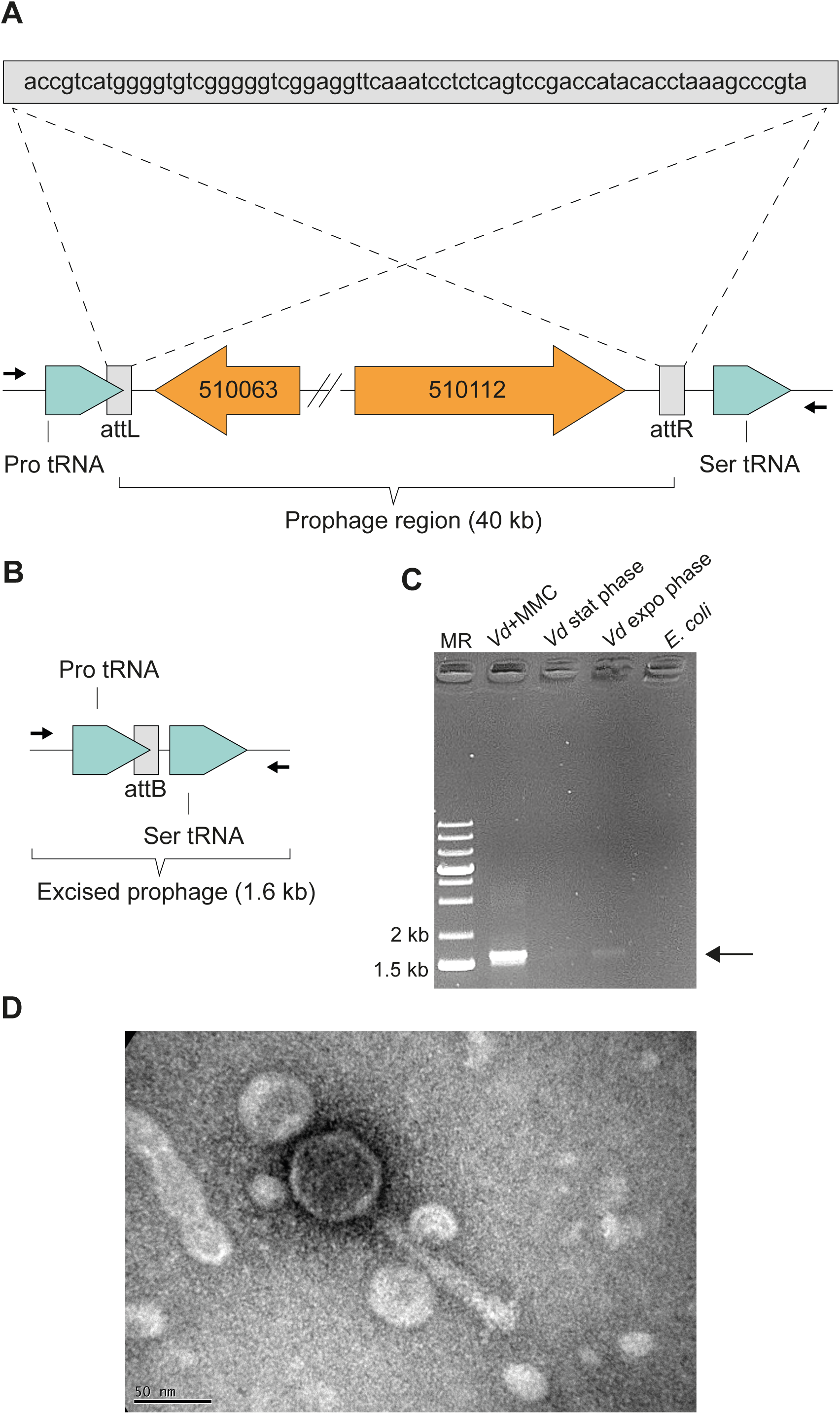
Genomic region of Vdi_1 and prophage excision. (A) Localization of the insertion site of Vdi_1. Note the repeated sequences attL and attR flanking Vdi_1 and its insertion at the 3’-end of Pro tRNA. (B) Genomic region of *V. diazotrophicus* NS1 upon prophage excision. Arrows represent the primers used to test Vdi_1 excision. (C) electrophoresis gel showing prophage excision, using primers flanking Vdi_1 prophage. MMC: Mitomycin C. (D) Transmission Electron Microscopy picture of Vdi_1 particle, released upon spontaneous excision from *V. diazotrophicus* NS1 grown in LB.

Importantly, Vdi_1 constitutes the only prophage region detected by Phigaro, suggesting that *V. diazotrophicus* NS1 is endowed with only one prophage. Moreover, Vdi_1 prophage DNA was not detected in the genome of 8 other *V. diazotrophicus* strains (strains 60.18M, 65.7M, 60.27F, 60.6F, 65.10M, 99A, 60.6B and HF9B, data not shown) and its closest relative is a prophage from *Vibrio plantisponsor* LMG24470 (**Fig. S4**).

### Vdi_1 is an active prophage

We next sought to demonstrate that Vdi_1 is an active prophage. If this hypothesis proves true, the viral DNA should excise by homologous recombination between the *attL* and *attR* sites of the prophage region (**Fig. 3A**). Viral DNA excision should therefore lead to the removal of the 40 kb prophage region (**Fig. 3B**). A PCR was conducted, using a primer pair annealing upstream and downstream of the *attL* and *attR* sites, respectively. Successful amplification of a 1.6 kb fragment would demonstrate successful excision of a prophage (**Fig. 3B**). We used *V. diazotrophicus* genomic DNA as template, extracted from 3 growing conditions: LB in exponential phase, LB in stationary phase, LB + Mitomycin C (MMC), a known prophage inducer. Indeed, a bright band at 1.6 kb was observed when using the DNA sample from MMC-induced culture, indicating massive 40 kb-loss in this condition (**Fig. 3C**). This experiment proved that Vdi_1 prophage is indeed an active prophage, being able to excise from the genome upon induction. Moreover, a faint-but-visible band was observed when using uninduced DNA samples (both from exponential and stationary phase cultures), indicating that Vdi_1 was able to activate and excise spontaneously (**Fig. 3C**).

Finally, a LB-grown overnight culture was centrifugated and the supernatant was ultracentrifugated to concentrate potential viral particles. The resuspended pellet was observed by transmission electron microscopy, which revealed the presence of intact phage particles, myovirus-type characterized by an icosahedral head, a thick tail with visible tail fibres (**Fig. 3D**). The presence of phage particles, together with the fact that the Vdi_1 prophage region was the sole region detected in the genome of *V. diazotrophicus* NS1 by Phigaro, formally demonstrated that Vdi_1 is indeed an active prophage.

### Generation of a prophage-free mutant

In order to understand the potential role(s) of Vdi_1 in the physiology of *V. diazotrophicus* NS1, we decided to create a prophage-free deletion mutant of this strain by deleting the entire Vdi_1 40 kb region using a dedicated suicide plasmid After two recombination steps, PCR amplifications were performed on potential mutants. The first primer pair flanked the entire Vdi_1 region, and gave a bright 1.6 kb band in uninduced cells, revealing the 40-kb deletion. However, as spontaneous excision can occur (**Fig. 3C**), we used a second primer pair, using primers amplifying a fragment within the Vdi_1 region. The presence of an amplification band in the wild type and its absence in the mutant confirmed that a Vdi_1 deletion mutant was obtained. Finally, we were unable to observe any phage particle in the supernatant of the mutant, unlike the one of the wild type (data not shown) supporting that Vdi_1 was the only prophage of *V. diazotrophicus* NS1 and that the Vdi_1-deletion mutant corresponded to a prophage-free mutant.

### A prophage-free V. diazotrophicus mutant is more resistant to mitomycin induction

In a second set of experiments, *V. diazotrophicus* NS1 and its prophage-free derivative mutant were grown in LB under oxic conditions. Mitomycin C (MMC) was spiked at different concentrations, and OD_600nm_ was monitored regularly over 10hour period to track biomass concentrations. A drastic decrease in OD was observed for *V diazotrophicus* NS1 with increasing MMC concentrations, dropping from OD 2.4 without MMC to OD 0.5 at 0.625 µg.ml^-1^ MMC (**Fig. 4A**). A same trend was observed for the prophage-free *V. diazotrophicus* mutant, confirming that MMC is a potent anti-microbial (21). Interestingly, at intermediate MMC concentrations, the decrease in final biomass concentration (measured as OD600_nm_) largely differed between the wild type and the mutant, with a significantly greater decrease observed for the wild type. Indeed, growth curves largely overlap between both strains, and start to diverge around 200 minutes post-MMC spiking (**Fig. 4**, **inlet**). This difference is likely due to the induction of Vdi_1 in the wild type strain, causing cell lysis and faster OD decrease. Supporting this hypothesis, timelapse movies, acquiring upon MMC induction at 0.0125 µg.ml^-1^, showed frequent and rapid cell elongation with aberrant cell shape in wild type cells, an effect not observed in the prophage-free mutant (**supplementary video S1** and **S2**).

**Fig. 4.**
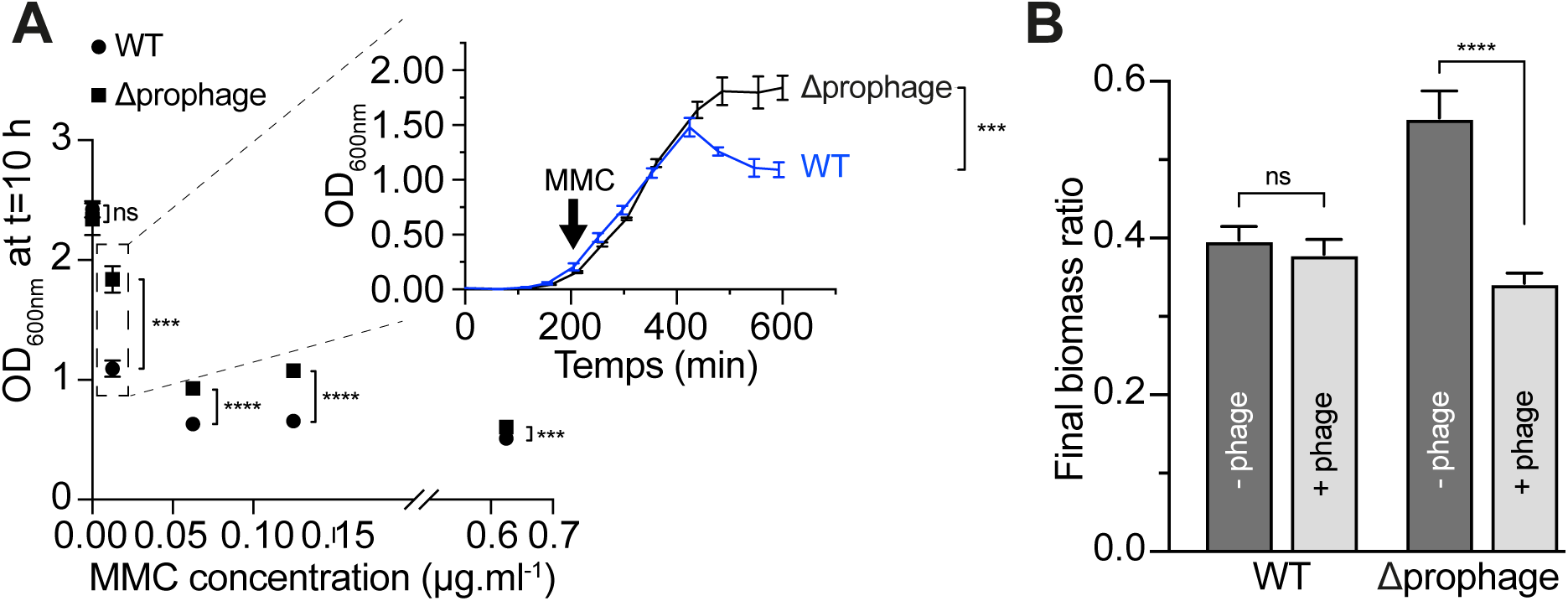
Prophage activation and infection experiments. Sensitivity of *V. diazotrophicus* NS1 and the prophage-free mutant to MMC (A). Cultures of both strains were inoculated with MMC at increasing concentrations, and the OD_600nm_ measured after 10 hours. An unpaired two-tailed t-test for every MMC concentration was performed. ns=not significant. Inlet, entire growth curves for both strains when grown in LB supplemented with 0.0125 µg/ml MMC. Infection experiment using Vdi_1 and *V. diazotrophicus* NS1 or its prophage-free derivative as preys (B). Vdi_1 particles (for the “+Phage” histograms) and the supernatant of the prophage-free mutant (“-Phage” histograms) were used to inoculate either *V. diazotrophicus* NS1 or the prophage-free mutant as prey. Cultures were incubated for 27 hours, and the final OD_600nm_ was measured. The y-axis corresponds to the ratio of the final OD_600nm_ obtained in the test experiment by the final OD_600nm_ obtained when adding the TPV-1 buffer. An unpaired two-tailed-test was done to compare the two conditions. ns=not significant.

### Vdi_1 produces infective particles

We then tested whether Vdi_1 particles are able to infect *V. diazotrophicus* cells. Phage particles were collected and used to infect either *V. diazotrophicus* NS1 or the prophage-free mutant. The mutant was included as prey because the prophage located in the genome of the wild type might confer innate resistance to secondary infection by the same virus. However, despite repeated attempts under multiple conditions (+/-O_2_, in LB or MDV media, using either non-induced or MMC-induced phage-producing cells, see material and methods), we were unable to obtain lysis plaques on plates, both using *V. diazotrophicus* or its prophage-free derivative mutant as prey (data not shown). We subsequently measured the growth of both strains, either spiked with the phage suspension or spiked with the suspension from the prophage-free mutant. The results showed that the presence of phage particles induced a significant decrease (p<0.0001) in the biomass in the prophage-free mutant as prey (**Fig. 4B**). This shows that, despite the absence of visible plaque lysis, Vdi_1 was able to infect and kill prophage-free mutant cells, leading to decreased final biomass.

### Vdi_1 does not play a role in N_2_-fixation under micro-oxic conditions

In order to determine the role Vdi_1 might have in *V. diazotrophicus*, we first sought to determine the impact of Vdi_1 deletion in the physiology of this strain. Our results first demonstrated that the deletion of the Vdi_1 prophage region did not modify the growth of *V. diazotrophicus* when grown in liquid LB, reinforcing the fact that Vdi_1 was generally silent in this strain or not able to infect the wild-type prophage-containing host (**Fig. S5A**).

Moreover, when both the wild type and the prophage-free mutant were grown in softgellan medium, the strain in which Vdi_1 region was deleted is still able to grow in the absence of nitrogen sources other than N_2_ and to fix N_2_ at a comparable rate as the wild type (**Fig. S5 BC**). Therefore, the prophage has no role in the diazotrophic activity of *V. diazotrophicus* NS1 when grown under micro-oxic conditions.

### Vdi_1 prophage participates in biofilm production in V. diazotrophicus

We recently demonstrated that *V. diazotrophicus* NS1 produces a thick biofilm, and hypothesized that this biofilm production is an ecophysiological adaptation of this strain to limit O_2_ diffusion with the biofilm, creating micro-oxic microenvironments suitable for diazotrophy (14). As biofilm matrixes are known to be composed of polysaccharides, but also of DNA, RNA and proteins, we hypothesized that Vdi_1-induced cell lysis leads to the release of those organic molecules, which participates in biofilm structuration. To test this hypothesis, we grew *V. diazotrophicus* NS1 and its derivative prophage-free mutant in microplates without shaking, using different growth media, under both oxic and anoxic conditions, and assessed biofilm production after 24 and 48 hours of growth. Under oxic conditions, no significant difference was observed between both strains when grown both under static condition (in microplate) and in milli-fluidic using Confocal Laser Scanning Microscopy (Fig. S6). In contrast, under anoxic conditions, the prophage-free mutant tended to produce less biofilm than the wild type, a trend which is statistically significant when supplemented with NH_4_^+^ (**Fig. 5**). These results highlighted that Vdi_1 may participate in biofilm production in anoxic environments, likely through phage activation and cell lysis.

**Fig. 5.**
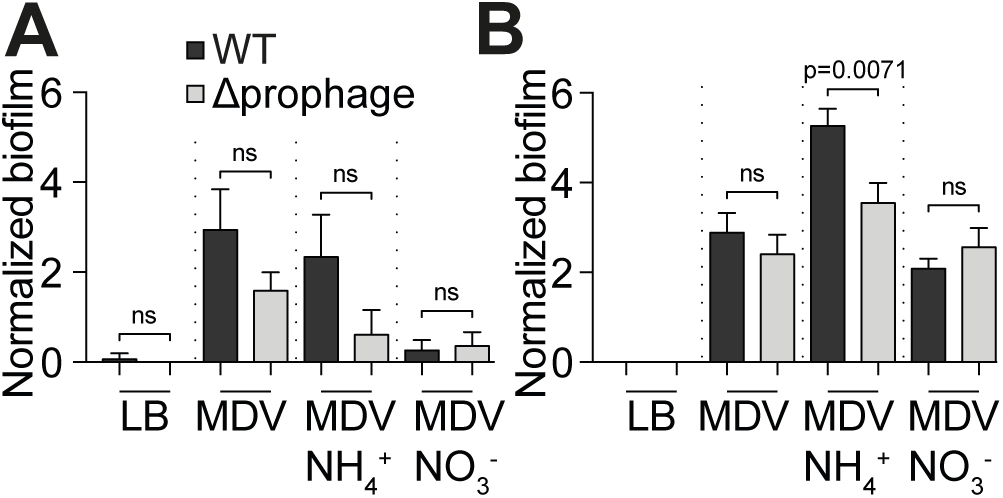
Biofilm production by *V. diazotrophicus* NS1 or the prophage-free mutant under anoxic conditions. Both strains were grown for 24 hours (A) or 48 hours (B) in microplates under anoxic conditions, in different media. Biofilm production was normalized by dividing OD_550nm_ by the OD_600nm_ obtained after 24 (A) or 48 h (B). Unpaired two-tailed t-tests were performed. ns=not significant.

### Vdi_1 prophage activation leads to the release of carbon and nitrogen compounds

As prophage induction leads to host cell death (**Fig. 4**), we hypothesized a release of C and N in the external environment during the phage-mediated cell lysis. To test this hypothesis, we first induced with MMC both the wild type strain and the prophage-free mutant. Then the amount and nature of C and N compounds in the filtrates were quantified and compared with non-induced samples. Dissolved Organic Carbon (DOC) was detected in MMC-free samples for both the wild type and the prophage-free mutant (4.5 to 5.8 mg.l^-1^ from 2x10^9^ cells, **Table S6**). MMC induction led to a 2.5-fold increase in DOC release in the wild-type strain, but only to a 1.4-fold increase in the prophage-free mutant, and this difference was significant (**Fig. 6A**). This increased DOC release was mostly due to the increased release of Low Molecular Weight (LMW) DOC (**Fig. 6B** and **Table S6**), while Hydrophobic DOC was released irrespective of MMC induction, and High Molecular Weight (HMG) and Humic DOC were only very modestly detected (**Table S6**). Strikingly, MMC induction also led to the release of urea in both strains, a molecule which was completely absent in uninduced ones (**Fig. 6C**). Moreover, ammonium releases significantly (*p* < 0.05) increased after induction in the wild type samples but not in the prophage-free mutant (**Fig. 6D**).

**Fig. 6.**
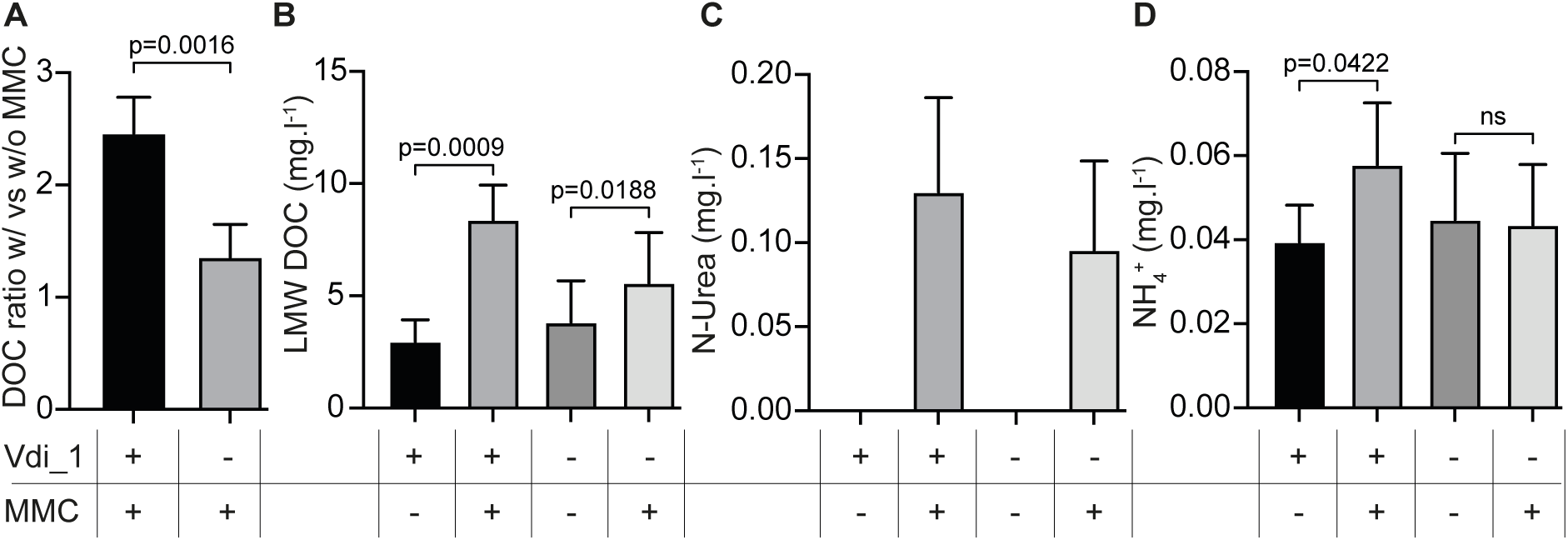
Carbon and nitrogen released by *V. diazotrophicus* and the prophage-free mutant. Total DOC amount released upon MMC induction divided by the total DOC amount released without induction. An unpaired two-tailed Welch’s t test was performed (A). Low Molecular Weight (LMW) (B), N-urea (C) and NH_4_^+^ (D) released by both strains without and with MMC induction. Paired two-tailed t-tests were performed for (B-D).

## Discussion

Viruses are the most abundant entities on earth and have long been viewed as dangerous entities, causing death and destruction at their vicinity. As a consequence, viruses play a fundamental ecological role by controlling the number of hosts. In addition, virus-mediated cell lysis can lead to the release of (i) (in-)organic matter participating in nutrient recycling and (ii) particulate organic matter which can serve new micro-environments. However, the potential ecological roles of viruses remain largely unexplored and, are therefore frequently overlooked in environmental studies. In this study, we provide evidences that a prophage of the marine diazotroph *V. diazotrophicus* NS1 plays a role in biofilm structuration and in the release of C and N, demonstrating that viruses can be key players in biogeochemical cycles of these two elements.

In the recent years, marine NCDs have been detected associated with animals (22, 23) or in low-O_2_ environments (*e.g.* in Oxygen Minimum Zones, OMZ (24)), supporting the idea that hypoxic or anoxic environments are conducive for nitrogenase functioning. In addition, their frequent presence within large particles suspended in the water column (11, 25), suggests a tendency to aggregate and produce biofilm as a strategy to limit O_2_ diffusion. Indeed, sinking particles, including marine snow and fecal pellets, contain a high C:N ratio exceeding the Redfield ratio and can be considered as hotspots for marine NCDs (26). We therefore conducted a global transcriptomic approach on *V. diazotrophicus* NS1, comparing their profile when grown under SRN-limited and micro-oxic conditions with growth in anoxia or in biofilm. We first demonstrated that this strain engages in massive gene transcription reprogramming upon growth in both of these conditions (**Fig. 1** and **Fig. S1**), highlighting that these environmental shifts correspond to major lifestyle changes for this strain.

Strikingly, the entire *nif* cluster was down-regulated under these conditions (**Fig. S2**). The down-expression of the nitrogenase is consistent with the inability of *V. diazotrophicus* NS1 to fix N_2_ under anoxic condition (12), while micro-oxic condition has been shown to be conducive to diazotrophy (14). Therefore, the observed decreased nitrogenase expression in anoxia shows that a low-but-not-null O_2_ tension is necessary for diazotrophy to occur, probably to produce ATP by respiration which in turn will fuel the nitrogenase. Enhanced nitrogenase activity under low O_2_ levels has been observed in some facultative anaerobe NCDs (13). while anoxia can also sustain diazotrophy in other NCDs (27, 28), highlighting the existence of various energetic strategies to fuel diazotrophy. The down-regulation of the *nif* cluster under biofilm condition is also unexpected and contradicts our previously published results highlighting an increased biofilm production under SRN limited conditions (14). This discrepancy may be due to the timing of RNA extraction for the RNAseq analysis, which provides only a snapshot image of possible successive events occurring during the time course of biofilm production.

In sharp contrast to the downregulation of the entire *nif* gene cluster, the RNAseq approach unveiled the existence of a single prophage within *V. diazotrophicus* NS1 that was upregulated both under anoxic and biofilm conditions. This prophage, coined Vdi_1 and belonging to family *Myoviridae*, is absent from related strains of the same species and its closest relative was found in the genome of *V. plantisponsor* LMG24470 (**Fig. S4**). We showed that Vdi_1 is indeed an active prophage, which (i) can spontaneously activate, (ii) be induced by MMC and (iii) form intact and infective phage particles (**Fig. 3-5**). Prophage activation typically occurs when host cells experience physiological stress such as low energetic status or oxidative stress, enabling the viruses to escape a possibly dying cell to find a new host. The activation of Vdi_1 under growth in anoxia or in biofilm suggests that such conditions trigger a stress to *V. diazotrophicus* NS1, probably linked to the energetic status of the host. This hypothesis is further supported by the overexpression of stress response genes like *ibpA* (BBJY01_360039, 40 to 47-fold in anoxia and biofilm, respectively, **Table S5**), encoding a small Heat-Shock Protein whose importance in biofilm formation has been demonstrated in *Vibrio fischeri* (29), and the CpxARP envelop stress system (BBJY01_490067 to 490069, induced up to 890-fold in anoxia, **Table S4**) and implicated in biofilm formation and resistance to stress in *Vibrio alginolyticus* (30). In the archaeon *Methanothermococcus thermolithotrophicus*, putative viral genes were shown to be overexpressed when grown under diazotrophic conditions, but no complete prophage was detected (28). To our knowledge, Vdi_1 represents the first member of demonstrated active prophage within marine NCDs. in providing fixed nitrogen to marine microbial communities, viruses that infect them likely represent a significant yet still underexplored component of marine ecosystem dynamics. Given the central role of diazotrophs in providing fixed nitrogen to the marine plankton community, diazotroph-infecting viruses represent an important-but-yet-understudied compartment of ecosystem functioning (31).

Numerous studies demonstrated prophage activation upon bacterial growth in biofilm (32), and this activation may participate in biofilm structuration (33, 34). Indeed, a bacterial biofilm is a complex structure composed of polysaccharides, but also of DNA, RNA and proteins (35). Here, we demonstrated that Vdi_1 prophage participates in biofilm formation in the marine diazotroph *V. diazotrophicus* NS1 under anoxic condition, probably through host-cell lysis, as a prophage-free mutant tended to produce less biofilm than the wild type (**Fig. 5**). Phage-mediated biofilm production can affect downstream processes, as shown in *Pseudomonas aeruginosa* carrying the prophage Pf4 which participates in the virulence of this strain (36), while a decreased virulence was observed in a phi458-deleted mutant strain of *E. coli* (37). As biofilm seems to be the preferred lifestyle of marine NCDs (11, 25), we suggest that prophage activation may regulate diazotrophy, and therefore oceanic biogeochemical cycles, by participating in biofilm structuration. On the other hand, we showed also that Vdi_1 has no role in nitrogenase activity when grown micro-oxically in softgellan, as a prophage-free mutant shows a similar nitrogenase activity as the wild type in this condition (**Fig. S5**). However, we recently demonstrated that, in this experimental context, cells are free living and do not form aggregates (14). Whether Vdi_1 indirectly affects diazotrophy when cells switch to multicellularity by biofilm production (*i.e.* when grown oxically under static condition) remains therefore to be demonstrated.

In addition to its role in biofilm production, we showed here that prophage-mediated host cell lysis triggers the release of significant amount of LMW-DOC (**Fig. 6A**) and ammonium. We also observed that Hydrophobic DOC is released without induction, whereas urea release is specifically induced by MMC-induced stress.. Therefore, it is tempting to speculate that *V. diazotrophicus* cells in a bad shape (*e.g.* because of low energetic status or oxidative stress) will undergo Vdi_1-induced cell lysis, which will lead to the release of SRN and carbon sources, to help clonemate survive in harsh conditions, through a process known as kin selection (38). Moreover, from an ecological point of view, the release of such molecules from diazotrophs may play a crucial role in ecosystem functioning, as the diazotrophically derived nitrogen can be used by the remaining (phyto-)planktonic community, fueling primary production. Interestingly, a recent RNAseq approach on *V. diazotrophicus* NS1 grown under diazotrophic conditions showed the strain’s capacity to assimilate both ammonium and urea, two N sources preferentially taken up by planktonic communities (12).

To our knowledge, the only report showing the role of a prophage in the release of fixed nitrogen came from *Trichodesmium* (39). Here, we demonstrated that Vdi_1-mediated cell lysis leads to the release of 6 µmol of DOC and 16.3 µmol of NH₄⁺ from 2x10^9^ cells (after subtracting the amount of DOC or NH_4_^+^ released by the prophage-free mutant upon MMC induction). This corresponds to a release of 3 fmol DOC and 8.15 fmol NH_4_^+^ per *V. diazotrophicus* cell. Assuming an abundance of 10^5^ Single-cell free-living NCDs per liter in surface waters (40). and given that most bacteria are endowed with at least one prophage, this corresponds to a potential phage-induced release of 300 pmol DOC and 815 pmol NH_4_^+^ per liter of seawater by NCDs. This potential release is highly conservative, as NCDs may grow at higher density when associated with aggregates and in aphotic waters or deep-sea sediments. Therefore, given the high prevalence of marine NCDs in ocean (11, 41, 42), phage-mediated host killing has the potential to significantly contribute to oceanic nitrogen and carbon cycle, especially in oligotrophic tropical and subtropical waters.

In conclusion, this study unveils the existence of a prophage in the marine NCD *V. diazotrophicus*. Its activation leads to biofilm structuration which is known to be hotspots for diazotrophy and to diazotroph-host-cell lysis leading to the release of SRN and carbon compounds to the environment. Given the prevalence of proviruses in bacterial and archaeal genomes including diazotrophs, our study provide evidences that proviruses may largely contribute to biogeochemical cycles and oceanic ecosystem functioning.

## Material and methods

### Strains and culture conditions

The *V. diazotrophicus* strain used in this study was *V. diazotrophicus* NBRC 103148 (also named NS1). *V. diazotrophicus* and *Escherichia coli* strains were routinely grown in Lysogeny Broth (LB Lennox) at 30°C and 37°C, respectively. When required, *V. diazotrophicus* was grown in SRN-deficient media (MDV for Modified Diazotrophic medium for *Vibrio* (12)). If necessary the following compounds were added to the media, either before autoclaving for NH_4_Cl (1 g.l^-1^), sodium thioglycolate (ref T0632, Sigma, 76 mg.l^-1^), resazurin (R7017, Sigma-Aldrich) (500 μg.l^-1^ from a 500 mg.l^-1^ stock solution), and agar (1.5%), or after autoclaving for cystein-HCl (ref 23255.186, VWR, 0.05% from a 5% filtered-sterilized solution in MQ water), NaNO_3_ (46,67 mg.l^-1^ from a 46,67 g.l^-1^ autoclaved stock solution), trimethoprim (Trim, 10 μg.ml^-1^), chloramphenicol (Cm, 5 μg.ml^-1^), glucose (Glc, 0.3 g.l^-1^), L-arabinose (L-ara, 0.2%), mitomycin C ((MMC, ref 51854, Ozyme) prepared at 5 mg.ml^-1^ in DMSO and diluted at 5 to 500 μg.ml^-1^ stock solutions in MQ water, used at concentrations between 0.001 and 1 μg.ml^-1^), DiAminoPimelic Acid (DAP, 0.3 mM).

Doubling times were determined by spectrophotometry by inoculating (in triplicate) overnight LB-grown cultures in 20 ml-containing 100 ml erlenmeyers with shaking, and by regularly measuring OD_600nm_. The doubling time was measured from the exponential growth phase of each culture. Strains, plasmids and primers used in this study are summarized in **Tables S1** to **S3**.

### Strain constructions and DNA techniques

Standard procedures were used for all molecular experiments, following the suppliers recommendations.

Prophage deletion was performed in *V. diazotrophicus* using an already established protocol (14, 43). Briefly, around fragments of 800 bp flanking the prophage region were PCR-amplified, fused together and cloned in the suicide plasmid pLP12 (44) using *Eco*RI and *Xma*I. The resulting pFD156 plasmid was introduced in *E. coli* β3914 (45), which served as a donor during bacterial conjugation with *V. diazotrophicus*. After plating on LB+Cm+Glc, obtained colonies in which the pFD156 plasmid integrated the genome of *V. diazotrophicus* were grown overnight in LB+L-ara followed by plating on LB+L-ara. Deletion of the prophage was verified by colony-PCR, using a primer pair flanking the prophage region (primers 240217 and 240218) and an internal primer pair (primers 240228 and 240230). Strain *V. diazotrophicus* in which the prophage has been deleted was stored in -80°C (strain 588). The prophage-free mutant, as well as *V. diazotrophicus* wild type, were stored at UBOCC culture collection, under the number UBOCC-M-3575 and number UBOCC-M-3576, respectively.

The *in silico* plasmid map and DNA sequences are available upon request.

### Soft-gellan assay

The ability of *V. diazotrophicus* and the prophage-free mutant to grow under SRN-free conditions was assess using a recently described soft-gellan assay (14), with the following modifications to obtained cleaner bands and results: instead of using an overnight culture of *V. diazotrophicus* strains, overnight cultures were back diluted in fresh LB, allowed to grow until OD_600nm_ reaches around 0.2-0.5, and 1 ml of these cultures was washed 3 times with MDV without yeast extract and adjusted to OD_600nm_ 0.5. 50 μl of these washed-cultures were used to inoculate soft-gellan tubes. Growth was demonstrated by the presence of a clear growing ring in the tube after 3 to 4 days of growth.

### Acetylene Reduction Assay

Strains were grown in soft gellan as described above, except that 100 μl of overnight culture, washed 3 times and resuspended in half the initial volume, were inoculated. Tubes were sealed with rubber stoppers with reversible edge (ref ZZ124591, Sigma-Aldrich). A 1 ml-gas-tight syringe (Hamilton) was used to remove 900 μl of the headspace of the tube. 900 μl of acetylene were subsequently injected. Tubes were placed at 20°C without shaking. Acetylene reduction to Ethylene by the nitrogenase was measured at day 3 and day 6 by gas chromatography (column: GS-Alumina (1153552, Agilent), 50 m x 0.53 mm, 0.25 μm : system (HP 6890 Series, Agilent). At each sampling point, the population-wide Ethylene to Acetylene ratio was quantified. Non-inoculated soft-gellan tubes served as negative controls, where only acetylene was detected.

### Biofilm production in microplates

The propensity of *V. diazotrophicus* and the prophage-free mutant to produce biofilm was quantified under various conditions, using crystal violet as recently presented (14). Briefly, *V. diazotrophicus* or the prophage-free mutant were grown overnight in LB, before being washed 3 times either with fresh LB or with MDV. Cultures were adjusted to OD_600nm_ 0.01 in LB, MDV, MDV+NO_3_^-^ or MDV+NH_4_Cl. 200 μl of these OD-adjusted cultures were inoculated in quadruplicates in polystyrene 96-well microplates (ref 330035, Dutscher). Microplates were placed for 24 or 48 hours at 30°C without shaking. At the end of the incubation, OD_600nm_ was measured in a microplate reader (TECAN Infinite M1000) to infer cell growth. Cell suspensions were removed by inversion, 220 μl of crystal violet (0.1%) were added into the wells, followed by 10 minutes incubation. After 2 rinsing steps (by plunging the microplate in a distilled water bath), microplates were dried for 24-48 hours at room temperature, upside down. Crystal violet was subsequently dissolved with 30% acetic acid for 10 minutes, before OD_550nm_ measurement. Biofilm production was corrected by dividing the OD_550nm_ measured by the OD_600nm_ corresponding to cell growth after 24 or 48 hours. For the quantification of biofilm production under anoxic condition, inoculated microplates were placed in GazPak EZ anaerobe pouch system (ref 260683, Grosseron), and processed as described above.

### Biofilm culture

*V. diazotrophicus* biofilms were grown at 30°C under hydrodynamic conditions in a three-channel flow cell (1 x 4 x 44 mm; Biocentrum, DTU, Denmark; (46)). The flow system was assembled, prepared and sterilized as described by (47). The substratum consisted of a microscope glass coverslip (24 x 50 mm; Knittel Glasser, Braunschweig, Germany). Each channel was inoculated with 250 µl of an overnight culture of *V. diazotrophicus* diluted to an OD_600_ of 0.1 in MD medium. A 2 h-attachment step was performed without any flow of medium. Then a 5 ml.h^-1^ flow of MDV medium was applied for 24 h using a Watson Marlow 205U peristaltic pump (Watson Marlow, Falmouth, UK).

Biofilms formed by *V. diazotrophicus* carrying the pFD086 were observed by monitoring the GFP fluorescence with a LSM 900 Confocal Laser Scanning Microscope (Zeiss, Oberkochen, Germany) using a 40x oil immersion objective. GFP was excited at 488 nm and fluorescence emission was detected between 500 and 550 nm. Images were acquired at intervals of 1 nm throughout the whole depth of the biofilm. ZEN 2.1 software (Zeiss, Oberkochen, Germany) and ImarisViewer (https://imaris.oxinst.com/imaris-viewer) were used for visualization and image processing. Quantitative analyses of image stacks were performed using COMSTAT software (http://www.imageanalysis.dk/) (48). At least three image stacks from each of three independent experiments were analyzed.

### RNA extraction

Liquid MDV: Overnight LB-grown preculture were washed 3 times with MDV without yeast extract. 200 µl of the washed preculture was inoculated in Erlenmeyer flasks (in quadruplicates) with 20 ml MDV and incubated with shaking (200 rpm) at 30°C for 24 h. Samples (5.10^8^ cells) were processed using the Direct-zol DNA/RNA miniprep kit (ref R2080, Zymo Research).

MDV under anaerobic condition: Overnight LB-grown preculture were washed 3 times with MDV without yeast extract. 200 µl of the washed preculture were inoculated (in quadruplicates) in 100 ml-containing 100 ml-vials (ref 33110, Sigma Aldrich)) sealed with a blue rubber stopper (ref 2048-11800, Bellco Glass). O_2_ was removed by adding sodium thioglycolate in the solution 2 of the MDV medium before autoclaving, by adding 1 ml Cystein-HCl 5% in the reconstituted MDV medium after autoclaving and by flushing the headspace for 2 minutes with N_2_ gas immediately after autoclaving, when the temperature was above 70°C (49). Absence of O_2_ was observed the day after, thanks to the addition of resazurin in solution 2 before autoclaving. Incubation was done for 24 hours at 30°C with shaking. Samples (5.10^8^ cells) were processed using the Direct-zol DNA/RNA miniprep kit.

MDV biofilm: Overnight LB-grown preculture were washed 3 times with MDV without yeast extract. 2 ml of this washed preculture were adjusted to OD_600nm_ of 0.01 with MDV with yeast extract, placed in a 24-well polystyrene microplate (ref CC77672-7524, Starlab). The microplate was incubated for 48 hours at 30°C without shaking. Cell suspension was removed by inversion. 150 µl of Tri Reagent (ref 2050-1-50, Zymo Research) were subsequently added and left for 1 hour at room temperature. To have enough material, 2 wells were pooled to form one replicate, and each quadruplicate were centrifuged at 12,000 g for 1 minute, before transferring the supernatant in a RNAse-and DNase-free Eppendorf tube. Samples were processed using the Direct-zol DNA/RNA miniprep kit, starting from step 2.

Every extracted RNA was deposited on gel and stored at -80°C until processing.

### RNA-sequencing and Data Analysis

Total RNA were sent to Eurofins (Germany). The company did the ribosomal RNA depletion, the DNase treatment, cDNA library preparation and Illumina sequencing (NovaSeq 6000 S4 PE150 XP). Sequencing was performed on 3 replicates per condition (the replicate with the lowest RIN (RNA Integrity Number) according to Eurofins standards was removed).

Initial bioinformatic analysis was performed by Eurofins: 22 millions of raw reads were obtained per sample, which were processed using RiboDetector (50) to remove reads classified as rRNA. High quality reads were obtained using fastp (51). At least 20,09 millions of “High Quality” reads per sample were obtained. Those reads were aligned to the reference genome of *V. diazotrophicus* (GCA_000740015.1) using STAR (52). Genome wise quantification was achieved using featureCounts (53). Differential Gene expression between 2 conditions was performed using R/Bioconductor package edgeR (54).

Differentially Expressed Genes (DEGs) were initially determined by Eurofins with a cutoff of 0.1 for adjusted p-values. We subsequently trimmed the results to retain DEGs having an adjusted p-value below 0.05, which was further corrected (False Discovery Rate (FDR)) using the Benjamini-Hochberg method (**Table S4**).

Figures presenting RNAseq data were obtained using an in-house written Python script.

The RNAseq data obtained in this study are available through NCBI under BioProject PRJNA1133960.

### Prophage bioinformatic analysis

The initial screening for phage detection was performed in the MicroScope platform (https://mage.genoscope.cns.fr/microscope/home/index.php) (55) using the Phigaro tool. This region was enlarged to a 40 kb-region (see results section) to consider it as a potential prophage region and the corresponding sequence was used for further analyses. Prophage genome analysis was performed with PhageScope (19), and Virfam (56) was used to classify the phage.

### Electronic microscopy

Precultures of *V. diazotrophicus* and the prophage-free mutant were inoculated in LB. These overnight cultures were used to inoculate 50 ml LB, which were incubated overnight at 30°C. Cultures were subsequently centrifuged at 8000 rpm for 15 minutes to remove bacterial cells. Supernatants were ultracentrifuged (Beckman Optima LE-80 K, 45Ti rotor) at 41 000 rpm for 90 minutes at 10°C. Pellets were resuspended with 150 µl of a dedicated viral buffer (Tris-HCl 10 mM, NaCl 100 mM, CaCl_2_ 5 mM). 5 µl were deposited on a TEM grid, 5 µl of uranyl acetate 2% were added before microscopic observation.

### Plaque assay

Phage particles from uninduced cultures were obtained as follows: a *V. diazotrophicus* 8 h preculture was inoculated in 100 ml-containing 500 ml LB Erlenmeyer, which was allowed to grow overnight before phage concentration by ultracentrifugation at 41,000 rpm during 90 minutes at 10°C (Beckman Coulter XE-90 ultracentrifuge, rotor : SW41-Ti, tube : 13,2 ml Ultra-Clear^TM^). The supernatant was removed and the viral pellet resuspended in 1 ml of viral buffer.

Phage particles from induced cultures were obtained by inoculating an overnight preculture in 100 ml LB. When OD_600nm_ reached 0.2, mitomycin C (MMC, 0.0125 or 0.125 µg.ml^-1^) was added 4 hours later, phage particles were collected as presented above.

100 µl of the viral suspension (undiluted or diluted ten times) were mixed with 100 µl of MDV-washed prey (either *V. diazotrophicus* or the prophage-free mutant, adjusted at OD_600nm_ =1 from a stationary-phase culture) and incubated for 30 minutes at 30°C. Subsequently, the mixture was used to inoculate 10 ml of an agar-top medium containing agar (7 g.l^-1^) with LB (20 g.l^-1^) supplemented with CaCl_2_, 2H_2_O (0.37 g.l^-1^ final concentration from a 147,01 g.l^-1^ stock solution), MgCl_2_ (0.24 g.l^-1^ final concentration from a 95.21 g.l^-1^ stock solution), before being poured on top of LB plates. When working under anoxic conditions, MMC (0.0125 µg.ml^-1^) was added in the agar top in which arabinose (2 g.l^-1^) was also added. Anoxia was obtained by placing plates in GazPack.

Plaque assays were also tested in MDV under anoxic conditions, during which MDV (instead of LB) was used in the agar-top medium and the mixture was poured on MDV plates, incubated in GazPacks.

The same experiments were performed in parallel with the supernatant of a prophage-free derivative mutant, which serves as a negative control (no phage production).

For every condition, plates were incubated for 24 h and 48 hours at 30°C before lecture.

### Infection assay in microplates

*V. diazotrophicus* and the prophage-free mutant were grown in LB, and phage suspension (or lack of it for the prophage-free mutant) was obtained as done as for the plaque assays. In parallel, overnight cultures of both strains were washed twice and diluted to OD_600nm_ of 0.01. 180 µl were deposited into polystyrene 96-well microplates, incubated in TECAN microplate reader at 30°C and OD_600nm_ was measured every 30 minutes. After 3 hours of growth, 20 µl of either viral buffer, phage suspension (coming from the wild type strain) or phage-free suspension (coming from the prophage-free mutant) were added to the wells, and OD_600nm_ was measured every 30 minutes for 24 hours. The final OD_600nm_ of each well (from the wild type strain or the mutant, in contact with either the phage suspension or the phage-free suspension) was divided by the mean of the final OD_600nm_ obtained from the corresponding inoculations spiked with the TPV-1 buffer.

### Timelapse microscopy

Overnight precultures of *V. diazotrophicus* or the prophage-free mutant were used to inoculate fresh LB. After 3 hours of re-growth, cells were deposited onto a 1% agarose patch containing MDV + MMC (0.0125 µg.ml^-1^). Images of the same field was taken every 30 minutes for 5 hours, using a Eclipse Ni-E microscope (Nikon) equipped with a 100X Apochromat oil objective.

### Quantification of carbon and nitrogen released

Both *V. diazotrophicus* and the prophage-free mutant were grown overnight in LB. The next morning, cultures were adjusted to OD_600nm_ 0.01 in Erlenmeyer containing 30 ml LB, in quintuplicate. After around 5 hours, OD_600nm_ were measured for each culture. Cells were then washed twice with 15 ml NaCl (35 g.l^-1^), and resuspended in 1 ml NaCl. Each cell suspension was used to inoculate 2 Erlenmeyer flasks (per suspension) each containing 15 ml NaCl (35 g.l^-1^) to achieve a final concentration of 2x10^9^ cells per flask (assuming OD_600nm_ 1 = 10^9^ cells.ml^-1^ in LB). In one of the 2 Erlenmeyer flasks, MMC was added (0.0125 µg.ml^-1^). After 3 hours of incubation at 30°C with shaking, the entire suspension was filtered using GF/F filters. The filtrate was recovered in vials, stored in -20°C until measurement. All materials used, including vials, filter holders, falcon tubes, forceps, syringes, Erlenmeyer flasks were acid-washed with 3HCl at a final concentration of 3.5%.

Few hours before measurements, sample were defrosted at 4°C in the dark. Dissolved organic matter (DOM) composition and size fractionation analysis was performed in filtrates using size exclusion chromatography with multi detectors (DOC-LABOR©) according to the methodology described in (57) and (58). Concentrations of bulk dissolved organic carbon (DOC), urea, ammonium and DOC content in the operationally defined fractions: hydrophobic DOC (non eluted), high molecular weight (HMW; > 10kDa), “humic-like” and low molecular weight compounds (LMW; < 0.5kDA), were quantified.

### Statistical analyses

All statistical analyses were performed using GraphPad Prism (version 8.0.1). Tests with a *p value* <0.05 were considered significant.

## Acknowledgments

The authors acknowledge the MicroScope platform (https://mage.genoscope.cns.fr/ microscope/home/index.php) for providing access to their platform. This work was supported by the “Rising Star” (Pays de la Loire Region), “DBM” (CNRS-INSB) and “EC2CO” (CNRS-INSU) grants, awarded to FD. PC was supported by the CNRS-MITI (GdR OMER). CG is the recipient of a doctoral fellowship (PhD BRIOCHE project) co-funded by the Université Bretagne Sud (UBS) and the Région Bretagne.

## References

1. Pi HW, Lin JJ, Chen CA, Wang PH, Chiang YR, Huang CC, Young CC, Li WH. 2022. Origin and evolution of nitrogen fixation in prokaryotes. Mol Biol Evol 39.

2. Rucker HR, Kacar B. 2024. Enigmatic evolution of microbial nitrogen fixation: insights from Earth’s past. Trends Microbiol 32:554–564.

3. Gallon JR. 1992. Reconciling the incompatible - N_2_ fixation and O_2_. New Phytol 122:571–609.

4. Flores E, Herrero A. 2010. Compartmentalized function through cell differentiation in filamentous *Cyanobacteria*. Nat Rev Microbiol 8:39–50.

5. Kelly MJ, Poole RK, Yates MG, Kennedy C. 1990. Cloning and mutagenesis of genes encoding the cytochrome bd terminal oxidase complex in *Azotobacter vinelandii*: mutants deficient in the cytochrome d complex are unable to fix nitrogen in air. J Bacteriol 172:6010–6019.

6. Poole RK, Hill S. 1997. Respiratory protection of nitrogenase activity in *Azotobacter vinelandii* - Roles of the terminal oxidases. Biosci Rep 17:303–317.

7. Cornejo-Castillo FM, Zehr JP. 2019. Hopanoid lipids may facilitate aerobic nitrogen fixation in the ocean. Proc Natl Acad Sci USA 116:18269–18271.

8. Carpenter EJ, Romans K. 1991. Major role of the cyanobacterium *Trichodesmium* in nutrient cycling in the north atlantic ocean. Science 254:1356–8.

9. Farnelid H, Turk-Kubo K, Ploug H, Ossolinski JE, Collins JR, van Mooy BAS, Zehr JP. 2019. Diverse diazotrophs are present on sinking particles in the North Pacific Subtropical Gyre. ISME J 13:170–182.

10. Leger-Pigout M, Navarro E, Menard F, Ruitton S, Le Loc’h F, Guasco S, Munaron JM, Thibault D, Changeux T, Connan S, Stiger-Pouvreau V, Thibaut T, Michotey V. 2024. Predominant heterotrophic diazotrophic bacteria are involved in Sargassum proliferation in the Great Atlantic Sargassum Belt. ISME J 18.

11. Delmont TO, Pierella Karlusich JJ, Veseli I, Fuessel J, Eren AM, Foster RA, Bowler C, Wincker P, Pelletier E. 2022. Heterotrophic bacterial diazotrophs are more abundant than their cyanobacterial counterparts in metagenomes covering most of the sunlit ocean. ISME J 16:927–936.

12. Cretin P, Mahoudeau L, Joublin-Delavat A, Paulhan N, Labrune E, Verdon J, Louvet I, Maguer JF, Delavat F. 2025. High metabolic versatility and phenotypic heterogeneity in a marine non-cyanobacterial diazotroph. Curr Biol 35:2659–2671 e3.

13. Bentzon-Tilia M, Severin I, Hansen LH, Riemann L. 2015. Genomics and ecophysiology of heterotrophic nitrogen-fixing bacteria isolated from estuarine surface water. mBio 6:e00929.

14. Joublin-Delavat A, Touahri K, Cretin P, Morot A, Rodrigues S, Jesus B, Trigodet F, Delavat F. 2022. Genetic and physiological insights into the diazotrophic activity of a non-cyanobacterial marine diazotroph. Environ Microbiol 24:6510–6523.

15. Dixon R, Kahn D. 2004. Genetic regulation of biological nitrogen fixation. Nat Rev Microbiol 2:621–31.

16. Wilhelm SW, Suttle CA. 1999. Viruses and Nutrient Cycles in the Sea: Viruses play critical roles in the structure and function of aquatic food webs. BioScience 49:781–788.

17. Williams KP. 2002. Integration sites for genetic elements in prokaryotic tRNA and tmRNA genes: sublocation preference of integrase subfamilies. Nucleic Acids Res 30:866–75.

18. Delavat F, Miyazaki R, Carraro N, Pradervand N, van der Meer JR. 2017. The hidden life of integrative and conjugative elements. FEMS Microbiol Rev 41:512–537.

19. Wang RH, Yang S, Liu Z, Zhang Y, Wang X, Xu Z, Wang J, Li SC. 2024. PhageScope: a well-annotated bacteriophage database with automatic analyses and visualizations. Nucleic Acids Res 52:D756–D761.

20. Nobrega FL, Vlot M, de Jonge PA, Dreesens LL, Beaumont HJE, Lavigne R, Dutilh BE, Brouns SJJ. 2018. Targeting mechanisms of tailed bacteriophages. Nat Rev Microbiol 16:760–773.

21. Tomasz M. 1995. Mitomycin C: small, fast and deadly (but very selective). Chem Biol 2:575–9.

22. Guerinot ML, Patriquin DG. 1981. N_2_-fixing vibrios isolated from the gastrointestinal tract of sea urchins. Can J Microbiol 27:311–7.

23. Domin H, Zimmermann J, Taubenheim J, Fuentes Reyes G, Saueressig L, Prasse D, Höppner M, Schmitz RA, Hentschel U, Kaleta C, Fraune S. 2023. Sequential host-bacteria and bacteria-bacteria interactions determine the microbiome establishment of *Nematostella vectensis*. Microbiome 11.

24. Loescher CR, Grosskopf T, Desai FD, Gill D, Schunck H, Croot PL, Schlosser C, Neulinger SC, Pinnow N, Lavik G, Kuypers MM, LaRoche J, Schmitz RA. 2014. Facets of diazotrophy in the oxygen minimum zone waters off Peru. ISME J 8:2180–92.

25. Reeder CF, Filella A, Voznyuk A, Coet A, James RC, Rohrer T, White AE, Berline L, Grosso O, van Dijken G, Arrigo KR, Mills MM, Turk-Kubo KA, Benavides M. 2025. Unveiling the contribution of particle-associated non-cyanobacterial diazotrophs to N_2_ fixation in the upper mesopelagic North Pacific Gyre. Commun Biol 8:287.

26. Riemann L, Rahav E, Passow U, Grossart HP, de Beer D, Klawonn I, Eichner M, Benavides M, Bar-Zeev E. 2022. Planktonic aggregates as hotspots for heterotrophic diazotrophy: the plot thickens. Front Microbiol 13:875050.

27. Martinez-Perez C, Mohr W, Schwedt A, Durschlag J, Callbeck CM, Schunck H, Dekaezemacker J, Buckner CRT, Lavik G, Fuchs BM, Kuypers MMM. 2018. Metabolic versatility of a novel N_2_-fixing Alphaproteobacterium isolated from a marine oxygen minimum zone. Environ Microbiol 20:755–768.

28. Maslac N, Sidhu C, Teeling H, Wagner T. 2022. Comparative transcriptomics sheds light on remodeling of gene expression during diazotrophy in the thermophilic methanogen *Methanothermococcus thermolithotrophicus*. mBio 13:e0244322.

29. Chavez-Dozal A, Hogan D, Gorman C, Quintanal-Villalonga A, Nishiguchi MK. 2012. Multiple Vibrio fischeri genes are involved in biofilm formation and host colonization. FEMS Microbiol Ecol 81:562–73.

30. Zhang Y, Kang X, Wu F, Lu Y, Gan Z. 2025. The CpxA-CpxR two-component system regulates stress tolerance and virulence of *Vibrio alginolyticus*. Int J Biol Macromol doi:10.1016/j.ijbiomac.2025.144279:144279.

31. Glass JB, Rousk K. 2024. Microbial nitrogen transformation processes across environments: more than just a cycle. Trends Microbiol 32:519–521.

32. Pires DP, Melo LDR, Azeredo J. 2021. Understanding the complex phage-host interactions in biofilm communities. Annu Rev Virol 8:73–94.

33. Godeke J, Paul K, Lassak J, Thormann KM. 2011. Phage-induced lysis enhances biofilm formation in *Shewanella oneidensis* MR-1. ISME J 5:613–26.

34. Carrolo M, Frias MJ, Pinto FR, Melo-Cristino J, Ramirez M. 2010. Prophage spontaneous activation promotes DNA release enhancing biofilm formation in *Streptococcus pneumoniae*. PLoS One 5:e15678.

35. Karygianni L, Ren Z, Koo H, Thurnheer T. 2020. Biofilm matrixome: Extracellular components in structured microbial communities. Trends Microbiol 28:668–681.

36. Rice SA, Tan CH, Mikkelsen PJ, Kung V, Woo J, Tay M, Hauser A, McDougald D, Webb JS, Kjelleberg S. 2009. The biofilm life cycle and virulence of *Pseudomonas aeruginosa* are dependent on a filamentous prophage. ISME J 3:271–82.

37. Li D, Liang W, Hu Q, Ren J, Xue F, Liu Q, Tang F. 2022. The effect of a spontaneous induction prophage, phi458, on biofilm formation and virulence in avian pathogenic *Escherichia coli*. Front Microbiol 13:1049341.

38. Granato ET, Foster KR. 2020. The evolution of mass cell suicide in bacterial warfare. Curr Biol 30:2836–2843 e3.

39. Hewson I, Govil SR, Capone DG, Carpenter EJ, Fuhrman JA. 2004. Evidence of Trichodesmium viral lysis and potential significance for biogeochemical cycling in the oligotrophic ocean. Aquat Microb Ecol 36:1–8.

40. Pierella Karlusich JJ, Pelletier E, Lombard F, Carsique M, Dvorak E, Colin S, Picheral M, Cornejo-Castillo FM, Acinas SG, Pepperkok R, Karsenti E, de Vargas C, Wincker P, Bowler C, Foster RA. 2021. Global distribution patterns of marine nitrogen-fixers by imaging and molecular methods. Nat Commun 12:4160.

41. Turk-Kubo KA, Gradoville MR, Cheung S, Cornejo-Castillo FM, Harding KJ, Morando M, Mills M, Zehr JP. 2023. Non-cyanobacterial diazotrophs: global diversity, distribution, ecophysiology, and activity in marine waters. FEMS Microbiol Rev 47.

42. Delmont TO, Quince C, Shaiber A, Esen OC, Lee ST, Rappe MS, McLellan SL, Lucker S, Eren AM. 2018. Nitrogen-fixing populations of *Planctomycetes* and *Proteobacteria* are abundant in surface ocean metagenomes. Nat Microbiol 3:804–813.

43. Morot A, El Fekih S, Bidault A, Le Ferrand A, Jouault A, Kavousi J, Bazire A, Pichereau V, Dufour A, Paillard C, Delavat F. 2021. Virulence of *Vibrio harveyi* ORM4 towards the European abalone *Haliotis tuberculata* involves both quorum sensing and a type III secretion system. Environ Microbiol 23:5273–5288.

44. Luo P, He X, Liu Q, Hu C. 2015. Developing universal genetic tools for rapid and efficient deletion mutation in *Vibrio* species based on suicide T-vectors carrying a novel counterselectable marker, Vmi480. PLoS One 10:e0144465.

45. Le Roux F, Binesse J, Saulnier D, Mazel D. 2007. Construction of a *Vibrio splendidus* mutant lacking the metalloprotease gene *vsm* by use of a novel counterselectable suicide vector. Appl Environ Microbiol 73:777–84.

46. Pamp SJ, Sternberg C, Tolker-Nielsen T. 2009. Insight into the microbial multicellular lifestyle via flow-cell technology and confocal microscopy. Cytometry A 75:90–103.

47. Tolker-Nielsen T, Sternberg C. 2011. Growing and analyzing biofilms in flow chambers. Curr Protoc Microbiol Chapter 1:Unit 1B 2.

48. Heydorn A, Nielsen AT, Hentzer M, Sternberg C, Givskov M, Ersboll BK, Molin S. 2000. Quantification of biofilm structures by the novel computer program COMSTAT. Microbiol-Sgm 146:2395–2407.

49. Agranier E, Crétin P, Joublin-Delavat A, Veillard L, Touahri K, Delavat F. 2024. Development and utilization of new O_2_-independent bioreporters. Microbiol Spectr 12.

50. Deng ZL, Munch PC, Mreches R, McHardy AC. 2022. Rapid and accurate identification of ribosomal RNA sequences via deep learning. Nucleic Acids Res 50:e60.

51. Chen S, Zhou Y, Chen Y, Gu J. 2018. fastp: an ultra-fast all-in-one FASTQ preprocessor. Bioinformatics 34:i884–i890.

52. Dobin A, Davis CA, Schlesinger F, Drenkow J, Zaleski C, Jha S, Batut P, Chaisson M, Gingeras TR. 2013. STAR: ultrafast universal RNA-seq aligner. Bioinformatics 29:15–21.

53. Liao Y, Smyth GK, Shi W. 2014. featureCounts: an efficient general purpose program for assigning sequence reads to genomic features. Bioinformatics 30:923–30.

54. Robinson MD, McCarthy DJ, Smyth GK. 2010. edgeR: a Bioconductor package for differential expression analysis of digital gene expression data. Bioinformatics 26:139–40.

55. Vallenet D, Calteau A, Dubois M, Amours P, Bazin A, Beuvin M, Burlot L, Bussell X, Fouteau S, Gautreau G, Lajus A, Langlois J, Planel R, Roche D, Rollin J, Rouy Z, Sabatet V, Medigue C. 2020. MicroScope: an integrated platform for the annotation and exploration of microbial gene functions through genomic, pangenomic and metabolic comparative analysis. Nucleic Acids Res 48:D579–D589.

56. Lopes A, Tavares P, Petit MA, Guerois R, Zinn-Justin S. 2014. Automated classification of tailed bacteriophages according to their neck organization. BMC Genomics 15:1027.

57. Dulaquais G, Breitenstein J, Waeles M, Marsac R, Riso R. 2018. Measuring dissolved organic matter in estuarine and marine waters: size-exclusion chromatography with various detection methods. Environmental Chemistry 15:436–449.

58. Huber SA, Balz A, Abert M. 2011. New method for urea analysis in surface and tap waters with LC-OCD-OND (liquid chromatography–organic carbon detection–organic nitrogen detection). J Water Supply: Res Technol-Aqua 60:159–166.

